# Interpretable and Brain-Inspired Recurrent Model of Hierarchical Decision-Making Capturing Trial-by-Trial Variability

**DOI:** 10.1101/2025.11.13.688200

**Authors:** Amir.M Mousavi-Harris, Sajjad Zabbah, Jamal Esmaily, Reza Ebrahimpour

## Abstract

Humans often make decisions in hierarchical environments, where low-level perceptual judgments inform high-level strategies. Understanding how the brain computationally navigates such multi-level decisions remains an open challenge. While existing models offer statistical insights, they often fall short in capturing the neural mechanisms and trial-by-trial variability underlying individual choices. To address this gap, we developed a neurocomputational model that integrates a biologically inspired attractor network for low-level perceptual decisions with a recurrent neural network (RNN) for high-level strategic adjustments. With minimal architectural constraints, the RNN receives only raw firing rates, feedback, and prior environment, learning to infer environment-switching strategies without explicit access to confidence or stimulus strength. In a hierarchical task combining motion discrimination and bandit decisions (N = 9; ~10,800 trials), the model successfully reproduced three hallmark behavioral patterns observed in humans. Unlike previous models, the model also captured trial-to-trial variability in switching decisions and implicitly learned to estimate decision confidence. For interpretability, we used representational and sensitivity analyses. Representational analyses revealed internal dynamics consistent with evidence accumulation in the anterior cingulate cortex (ACC), while sensitivity analysis identified feedback as the dominant influence on strategy, modulated by recent trial history. This framework combines interpretability and predictive power, moving beyond simple data fitting to provide mechanistic insights into how the brain integrates confidence and feedback to guide adaptive behavior in hierarchical decision-making.

## Introduction

Hierarchical decision-making is a fundamental aspect of human behavior, encountered in everyday scenarios such as planning a vacation. At its core, hierarchical decision-making involves multiple interdependent layers of evaluation which makes the source of the error ambiguous. Consider a traveler choosing between a cultural city tour and a relaxing beach getaway. This high-level decision depends on various lower-level choices, such as selecting accommodations, assessing transportation reliability, and evaluating local attractions. If the overall experience is unsatisfactory (i.e. error), the source of this dissatisfaction is ambiguous, whether the destination itself did not meet expectations (i.e., an incorrect high-level choice) or whether the accommodation, transportation, or local attractions were poorly chosen (i.e., incorrect low-level decisions). It is still not well understood how the brain bridges different levels of hierarchy to resolve this ambiguity and shape higher-order strategies.

Empirical evidence from behavioral and neuroscientific studies, using controlled experiments, show that in both human and animal confidence in low-level choices serves as a proxy to solve the ambiguity of the error in hierarchical decisions^1–8^. If an agent is highly confident in their low-level decisions a negative outcome strongly suggests an error at the higher strategic level ^1,5,6^. Conversely, low confidence in low-level choices makes it difficult to pinpoint the exact source of the error. Research in human and nonhuman primates further reinforces these findings, demonstrating distinct neural correlates for different layers of decision-making ^5,9^. Sensory and association cortices encode low-level evaluations, while frontal regions—such as the anterior cingulate cortex and orbitofrontal cortex—are more involved in high-level strategic evaluation^5,10,11^.

Computational models show that, similar to low-order decision making process (e.g. drift-diffusion models ^18,19^, or recurrent dynamical systems ^22–24^), high-order decisions also rely on evidence accumulation over time until reaching a bound^14^–^17^. However, unlike low-order decisions where evidence is derived mainly from sensory inputs, evidence for high-order decisions is extracted from confidence in consecutive low-order choices. While such models provide statistical descriptions of neural and behavioral responses in hierarchical decision-making, they primarily account for average behavioral patterns and fail to capture complex dynamics, such as trial-by-trial variability and the nuanced neural signatures of high-level decisions.

Recently, recurrent neural networks (RNNs) have gained popularity for modeling complex neural dynamics due to their flexibility and ability to readily fit data via backpropagation ^25–29^. The flexible recurrent architecture usually let the model to capture the complex dynamics underlying higher-level decision processes. However, RNN solutions, while powerful, lack interpretability compared to traditional statistical and mechanistic approaches. To address this, researchers have employed various techniques, including representation analysis ^26^, fixed-point analysis ^29^, rank analysis ^30^, and constraints that align model behavior with neural data ^31^. Therefore, RNN-based models offer a framework that is sufficiently complex to capture trial-by-trial variability, while subsequent analyses can render these black-box models interpretable and mechanistically meaningful.

Here we propose an integrated approach that models low-level decision processes using traditional mechanistic models for interpretability, while employing RNN-based models at the higher level to capture complex decision dynamics. Specifically, we aim to model low-level trial by trial firing rates using an attractor neural network framework and integrate this with a high-level decision model based on overparameterized RNNs with LSTM cells. We hypothesize that RNNs can capture trial-by-trial variability in high-level decision-making by receiving output from a simulated neural firing rate. This allows us to probe the underlying neural dynamics in greater detail. Through techniques such as representational and sensitivity analyses, we will investigate how the model internally solves for hierarchical decision-making. Our approach not only captures the dynamics of low-level sensory processing and high-level strategic evaluation but also provides insight into the neural computations underlying hierarchical decisions, paving the way for future research in this domain.

## Methods and Materials

To keep it concise, we highlight the key elements of the methods here, with additional details available in the supplementary information.

### Participants

The study included nine participants, four men and five women, with an average age of 26 years (±5). Ethical approval for the experimental procedures was obtained from the Ethics Committee associated with the Iran University of Medical Science (Approval ID: IR.IUMS.REC.1400.1230). Importantly, all participants were unaware of the specific aims of the experiment and provided written consent after receiving a full briefing about the nature of the research. All participants either had normal vision or used corrective lenses to meet required standards. Participants underwent two data collection sessions across two days. Each session involved completing four sets of 150 trials, resulting in an average of 1,200 trials per participant and a total of 10,800 trials across all participants.

### Trial Procedure

Each trial began with participants fixing their gaze on a centrally located fixation point (FP). After a short delay, two bar targets appeared on opposite sides of the screen along with a random dot motion stimulus (see ^32^ for detailed task description). The main goal was to determine the overall direction of motion (either left or right) while also assessing confidence level. After a brief delay, the FP was disappeared, prompting participants to demonstrate their perceived direction of motion. Distinctive auditory tones were then used to provide feedback on the accuracy of their responses.

### Stimulus (Motion Direction Discrimination Training)

Each trial commenced with the subject fixating on a small red dot (0.3° diameter) located at the center of the screen. Following a variable delay period ranging from 200 to 500 milliseconds (governed by a truncated exponential distribution), two elongated bar targets (7° × 0.75°) appeared on opposite sides of the screen, equidistant from the fixation point at an eccentricity of 8°. A dynamic random dot stimulus was then presented within a circular aperture (5° diameter) centered on the fixation point for a duration of 500 milliseconds on each trial. Following the disappearance of the dynamic random dot stimulus, a variable delay period ranging from 400 to 1,000 milliseconds (also governed by a truncated exponential distribution) was introduced before the Go signal. The Go signal was indicated by the disappearance of the central fixation point. Subjects were instructed to maintain their gaze on the fixation point throughout the trial until the Go signal appeared. After receiving the Go signal, participants used a mouse to indicate their perceived direction of motion. They did this by moving the cursor to the corresponding target on the screen and clicking on it. To provide feedback, distinct auditory cues were used: a positive cue for correct responses and a negative cue for incorrect responses.

To assess participants’ confidence in their motion direction judgments, a color-coded confidence rating system was implemented. Confidence levels were quantified on a scale from 0 to 1, derived from raw mouse data. Specifically, participants indicated their confidence by clicking on a bar, and their response was normalized by dividing the click position by the total length of the bar, ensuring values ranged between 0 and 1. The system employed a visual color spectrum to represent confidence levels: green indicated the highest confidence (a value of 1), while red represented the lowest confidence (a value of 0). This intuitive design allowed participants to easily convey their certainty in their judgments.

Subjects underwent training on a basic motion discrimination task until achieving high performance, as evidenced by psychophysical thresholds (α, see ^14^) falling below 15%. To systematically manipulate task difficulty in the subsequent motion direction discrimination phase, random variations in motion strength were introduced across trials. Motion strength was defined by the percentage of dots exhibiting coherent movement and ranged from completely random (0%) to highly coherent (51.2%). Notably, for trials with 0% coherence, positive and negative feedback were delivered randomly with equal probability (50% each) to ensure that feedback was not inherently linked to a specific direction in these most challenging trials (Fig. S1).

### Hierarchical Decision-Making Task

After participants attained the criterion performance level, they proceeded to the changing environment task (Fig. 1A). The experimental setup, motion stimulus, and sequence of events remained consistent with those in the training phase. However, a modification was made regarding the number of choice bar targets presented. Instead of a single pair of choice bar targets, two pairs were now displayed—one above and one below the central fixation point— resulting in a total of four bar targets. These target sets were positioned 7° above and below the fixation point. Each pair corresponded to a distinct environment, with the left and right targets within each environment indicating possible motion directions.

**Figure 1.**
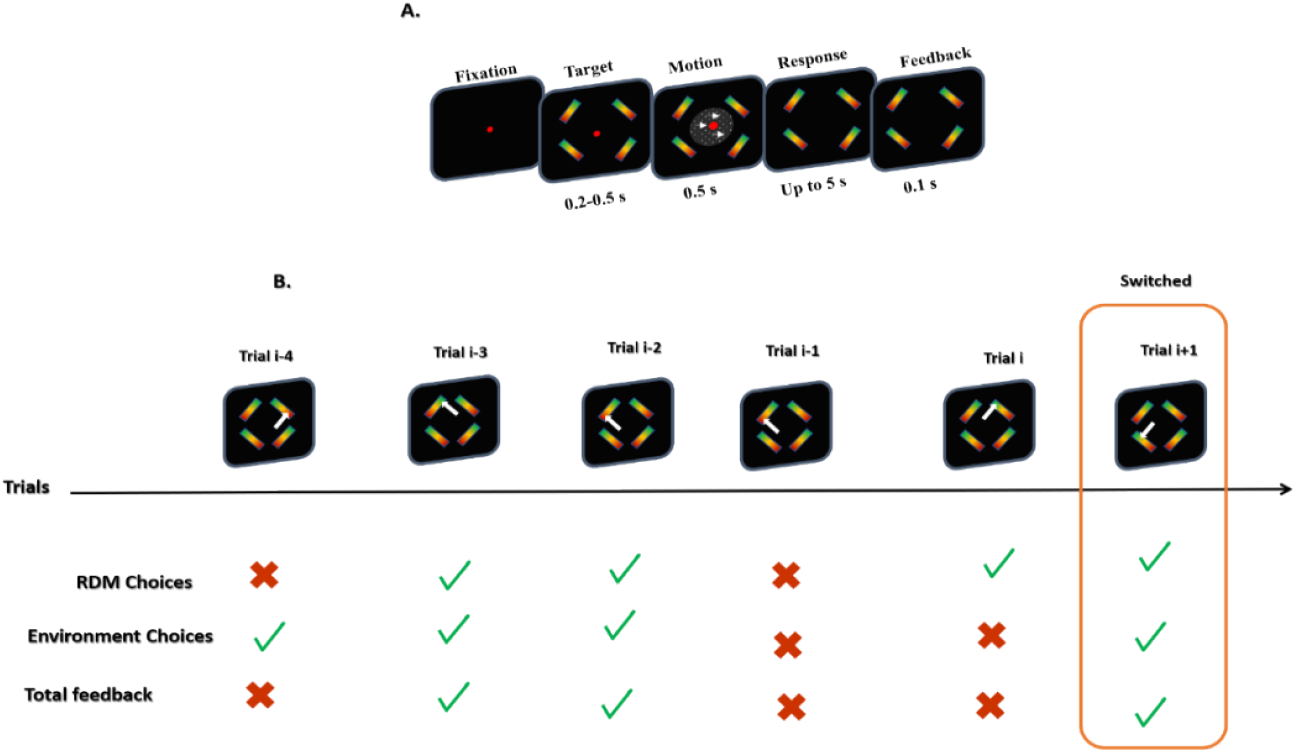
**(A) Task design**. Subjects were rewarded for correctly identifying both the appropriate environment and motion direction by selecting the corresponding targets. Each pair of targets, located above and below the fixation point (FP), represented different environments, with the right and left targets within each pair indicating possible motion directions. The rewarding environment remained unchanged for a variable number of trials (ranging from 2 to 15, drawn from a geometric distribution with a mean of 6) before switching without explicit notice. Throughout the trials, subjects had to infer the correct environment based on their history of feedback, choices, and confidence, despite random variations in motion direction and strength (%Coh – percentage of coherently moving dots). **(B) Example of Hierarchical Decision-Making**. This panel illustrates how subjects made decisions at two levels, along with their confidence, within a hierarchical structure. The upper panel shows the choices made during the task, while the lower panel displays the outcomes of both low-level and high-level decisions, as well as the feedback received. The white arrow indicates the subject’s choice.

When participants correctly identified the target matching both the correct environment and motion direction, they received positive feedback. If either or both choices were incorrect, negative feedback was given. This feedback was ambiguous, as it did not specify which of the choices was wrong. Participants simultaneously reported their confidence in the motion direction and made both direction and environment choices. The selection between left and right targets was termed the “direction choice,” while the selection between upper and lower targets was referred to as the “environment choice.” The active environment remained constant for a random number of trials before changing without any explicit cue.

### Neural Model

We introduced a neurocomputational architecture comprising two primary networks: the “Reasoning” module and the “Sensorimotor” module.

The Sensorimotor module involves the development of a neural attractor model based on previous work by Wong and Wang ^22,23^, which is a variation of the sequential sampling models for decision-making under uncertainty proposed by Bogacz et al. (2006) ^33^. These models have long been capable of accommodating various types of data ^34–36^. In this model, noisy sensory evidence is accumulated by two competing mechanisms (red and blue in Figure 2.A, left) that race toward a common predefined decision boundary while mutually inhibiting each other. The model makes a choice as soon as one mechanism reaches the boundary (for more information, see Supplementary Material). This model has accounted for numerous observations in both perceptual and value-based decision-making behavior, along with their underlying neuronal substrates in the human brain ^37^ and non-human primate brain ^38^.

**Figure 2.**
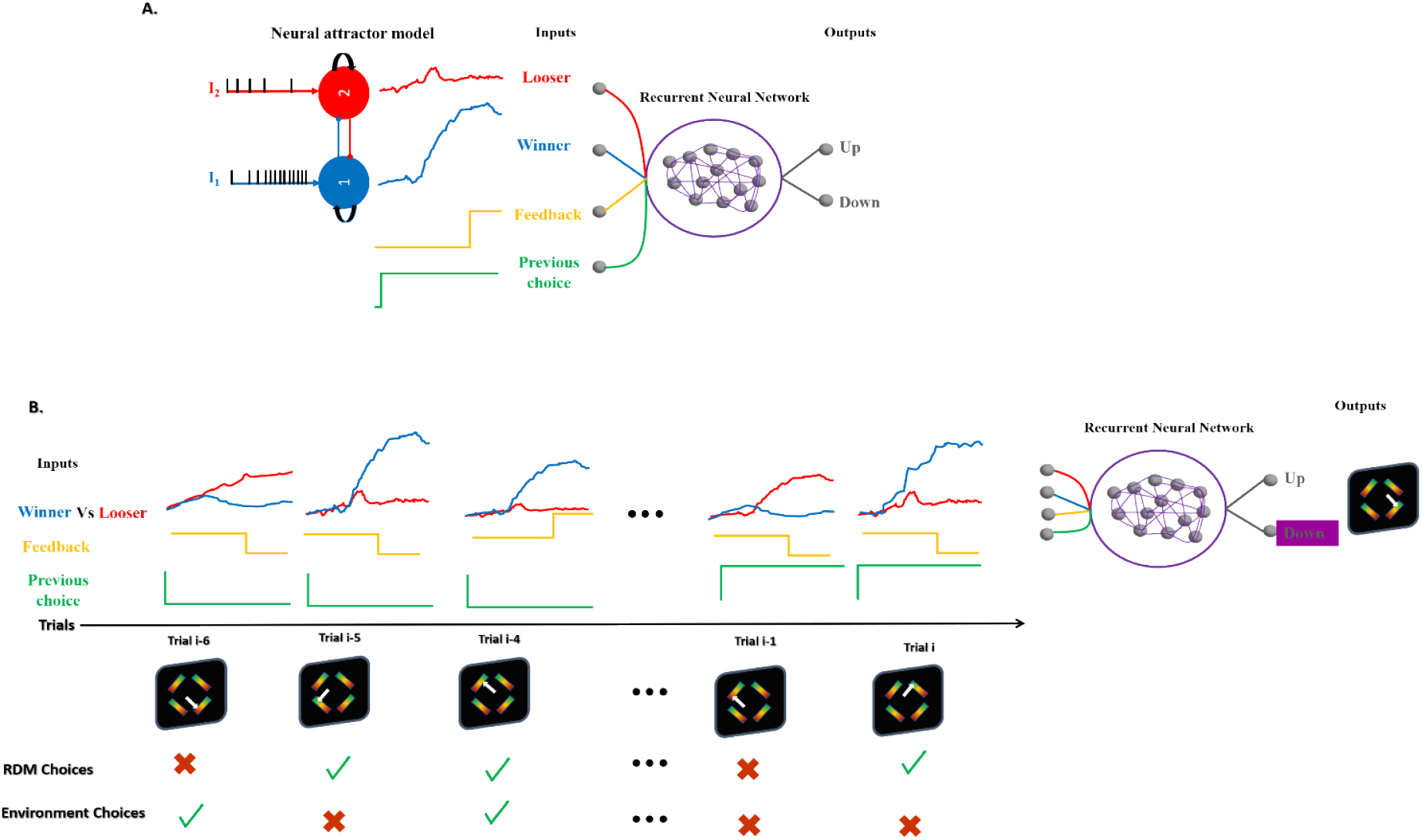
**(A) Schematic Overview of the model**. The high-level model consists of 256 LSTM neurons and leverages low-level decision signals (Loser: red, Winner: blue), feedback (yellow), and prior environment (green) to predict the next environment choice (“Up” or “Down”) via a single output unit. **(B) Schematic Overview of Training Data and Model Output**. This diagram illustrates the overall process of preparing training data and generating model output. Upper panel: Information from four trials is concatenated and fed into the model as a single input signal. The diagram shows four example signals, each with a length of 55 samples. Lower panel: This panel displays the subject’s choices and the associated confidence levels for each choice.

Model confidence was defined following previous work ^38–41^. It quantified the time-averaged difference between the activity of the winning and losing accumulators during stimulus presentation (0 to 500 ms), consistent with recent findings showing that confidence computations can continue after a decision is made, as long as sensory evidence remains available ^39,41–44^. More comprehensive details of the Sensorimotor module are provided in the section Sensory Module in the Supplementary Material.

### Parameters fitting

To calibrate the attractor network model and ensure its predictions closely matched participants’ behavior in isolated sessions, we performed a parameter fitting procedure focused on optimizing decision accuracy. The parameters adjusted during this process were the decision threshold (NDT) and the perceptual gain (µ_0_). The decision threshold determines the level of accumulated evidence required for the model to commit to a decision. This parameter directly influences the speed–accuracy trade-off: a lower threshold leads to faster but potentially less accurate decisions, while a higher threshold promotes more accurate but slower decisions. The perceptual gain scales the input current according to motion coherence, modulating the model’s sensitivity to the strength of sensory evidence.

The fitting procedure involved systematically varying NDT and µ_0_ across a predefined parameter space. For each parameter combination, the model was simulated over multiple trials designed to replicate the experimental task conditions. The model’s performance was evaluated by calculating its decision accuracy, defined as the proportion of trials in which the model selected the correct choice based on the coherence level. Our objective function was to minimize the discrepancy between the model’s output and participants’ empirical accuracy data. The parameter values that yielded the best decision accuracy were selected as the optimal fit for the model. The final optimized values for NDT and µ_0_ resulting from this fitting procedure are reported in Table S1. These optimized parameters allowed the model to successfully reproduce participants’ behavioral patterns in terms of choice accuracy across different task conditions.

### Hierarchical Module

To investigate the underlying mechanisms of high-level decision-making, we employed machine learning techniques to train the Reasoning module, implemented as a recurrent neural network (RNN) ^45^ with non-linear firing rate units. Long Short-Term Memory (LSTM) cells were used in the RNN architecture due to their ability to effectively model and retain long-term dependencies in sequential data. This architecture is particularly suited for dynamic problems like hierarchical decision-making, where information must be integrated over time.

We trained the network with minimal architectural constraints, focusing solely on achieving the desired outcome—solving the task ^9,29^. This allowed the network to discover its own solution strategy, offering valuable insights into the potential neural computations underlying human decision-making.

Designing the LSTM model required addressing two key questions:

1. What should be the input to the LSTM model?
2. How many trials, on average, are required to trigger an environment change?

Answering these questions was crucial for training the high-level neural network to effectively replicate the observed behavioral data. For the first question, through model simulations (Fig. S3) and building upon previous works ^5,6^, we found that feedback signals and the expected accuracy of low-level decisions were the critical elements influencing environment changes (Our simulations, detailed in Fig. S3, confirmed that other potential inputs did not yield effective performance). Therefore, we selected winning and losing signals from the low-level decision process (sensorimotor module), feedback signals, and the previous environment as inputs (see ^9^ and Fig. 2.A). The output of the network indicates an “Up” or “Down” choice for the next environment. Prior to this, we calibrated the sensorimotor module to provide the necessary input for the higher-level LSTM model.

To address the second question, we trained and evaluated the model’s performance using a range of window sizes, from 2 to 15. This specific range was chosen to align with the interval of environment changes observed in the experiment, as previously discussed (see Task Design description). Each window size was evaluated ten times, and the average performance across these runs is depicted in Figure S4.A. The results indicated that a window size of four emerged as the most effective configuration. Further details are provided in the Supplementary Materials (Fig. S4).

To optimize computational efficiency prior to model training and testing, we focused on reducing the signal length. Our initial data processing involved removing segments in which both winner and loser signals reached baseline (0–500 ms), as these intervals lacked informative content relevant to low-level decision-making. We then reduced the signal length from 1000 to 500 samples. However, this length remained computationally expensive. To address this, we employed a combined interpolation and averaging technique. Specifically, we implemented a fixed-size window that traversed the signal, calculating the average firing rate within each window. This process effectively reduced data dimensionality by first downsampling to 55 samples, and then further to 10 representative samples (Figure S4.B). Interestingly, training the model with this reduced data (three modes with lower dimensions) yielded results comparable to using the original 1000-sample data (Table S4). This finding indicates that the model can effectively capture essential information from the raw data, even with significant compression, suggesting that slow neural dynamics may be crucial for encoding high-level evidence.

### Training of LSTMs

Our main objective here was to train an LSTM model that replicates human data. Since we aimed to model the high-level decision-making process, we replicated the exact trajectory of participant behavioral data in the low-level decision-making. In this way, the low-level module was fixed, and only the high-level module was trained to produce the subjects’ high-level decision profile.

This low-level trajectory included coherence levels across trials, received feedback, degree of confidence, and the subject’s response (True/False). Due to the stochastic nature of the sensorimotor module, the low-level trajectory could be highly variable. To bridge this gap, the low-level model was run 500 times for each trial. The simulation that produced the smallest difference from the behavioral data—based on confidence, feedback, and overall accuracy—was selected to provide the most representative signals for the high-level model. Each feature was analyzed individually and separately. Notably, the closest match for each feature was determined using simple subtraction.

The four closest-matched signals were then fed into the high-level model using a fixed-size window: winner firing rate, loser firing rate, choice feedback, and the previous choice of environment. Notably, we did not provide the model with direct values for confidence, selection direction, or stimulus strength. Instead, we relied on the model’s ability to extract these crucial elements from the raw firing rate signals of the winner and loser accumulators.

It is important to note that the choice feedback signal remained at zero until the end of the stimulus presentation. Upon stimulus offset and decision announcement, the signal transitioned to either a positive or negative value, depending on the feedback outcome. An illustration of these input signals is provided in Figure 2.

To ensure that the network learned an optimal configuration of weights, we trained the LSTM model using TensorFlow ^46^. The training process employed two key techniques: backpropagation through time (BPTT) and the Adam optimizer ^47^.

The model parameters (weight matrices and bias vectors) were learned through a training process that minimized a chosen loss function (e.g., cross-entropy; Eq. 1) by iteratively updating the parameters using the backpropagation algorithm. The hyperparameters used in our LSTM model are presented in Table S2.

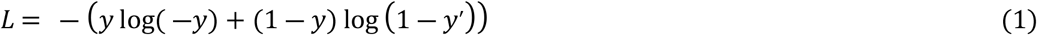

### Neural trajectory analysis

To investigate neural trajectories, we applied principal component analysis (PCA) to neural activity recorded across trials. The firing rate data used for this analysis had a shape of (number of trials, 220 time points, 256 neurons). Trials were selected based on experimental conditions, creating distinct trial groups. For each condition, we computed the mean neural activity by averaging across trials within each group, resulting in a mean activation pattern with a shape of (220 time points, 256 neurons).

To reduce dimensionality and visualize the underlying neural dynamics, we applied PCA separately to the mean activation pattern of each condition. PCA was performed across the 256 neurons, preserving the temporal dimension of 220 time points. For each condition, we extracted the top two principal components, which captured the greatest variance in the data. To illustrate the temporal evolution of neural activity, lines were drawn connecting the projected points of these top two principal components across the 220 time points.

To quantify how the model accumulates evidence, we computed the trajectory of the LSTM’s output and calculated the sum of the Euclidean distances from each trajectory point to the attractor point (absolute PCA values), representing the model’s final state. Additionally, we employed a bootstrap method with 1,000 iterations to estimate the variability of this measure. This sum was used to compare the model’s progression across different conditions, providing insights into how the LSTM’s internal dynamics evolve and accumulate evidence over time. The analysis was applied to each experimental condition to examine differences in how the model tracks and integrates information.

### Sensitivity analysis

To assess the feature importance of the input data, we utilized “saliency maps”, which allowed us to visualize the sensitivity of the model’s output with respect to changes in the input features. The model processed four features over time: Loser signal, Winner signal, Feedback, and Previous Environment. We computed the saliency map for each subject by calculating the gradient of the model’s output with respect to the input features. Specifically, we used TensorFlow’s GradientTape to track the gradients and compute the absolute values of the gradients relative to the input tensor.

For each subject, the saliency map was computed across all time steps and then averaged across subjects to assess general trends in feature importance. The mean saliency values were plotted along with their standard deviations over time, highlighting the relative contributions of different input features. To quantify the relative importance of each feature, we calculated the mean saliency values over time for each feature, along with the standard error of the mean (SEM) across subjects. These values were visualized using a bar plot, where the height of each bar represented the mean feature importance, and the error bars indicated the SEM.

Additionally, to investigate the recency effect in the saliency maps, we reshaped the mean saliency values to correspond to the four individual trials. This reshaped data allowed us to assess how feature importance evolved over time within and across trials. We computed the mean and SEM of saliency values across time steps for each feature in each trial. The resulting values were plotted in a bar chart with error bars, illustrating how feature importance changed across consecutive trials. This analysis provided insights into how different features influenced the model’s predictions over time, helping to interpret the role of each feature in the decision-making process and revealing recency effects in feature importance.

## Results

### Replication behavioral results

First and foremost, we verified that behavioral results, in both low-level and high-level decisions, successfully replicated key findings from Purcell and Kiani (2016) ^6^. Specifically, the interaction between low-level and high-level decisions was observed, validating the significance of three factors influencing high-level decisions: the number of negative feedbacks, stimulus strength, and trial persistence within an environment (Fig. 1, gray lines).

### The LSTM model extracts similar information as human participants to capture the probability of switching

We used the well-known attractor-based rate model to simulate participants’ behavior in response to perceptual stimuli (low-level model). This model accurately captures both performance and confidence trajectories as a function of stimulus strength (Fig. 3A and B). To account for trial-by-trial variability, we fine-tuned the noise in the low-level model by selecting the best noise vector for each trial using a blind search algorithm. This approach ensured that the simulated firing rates produced confidence levels most closely matching those of participants on individual trials, given specific stimulus strengths (Fig. S5).

**Figure 3.**
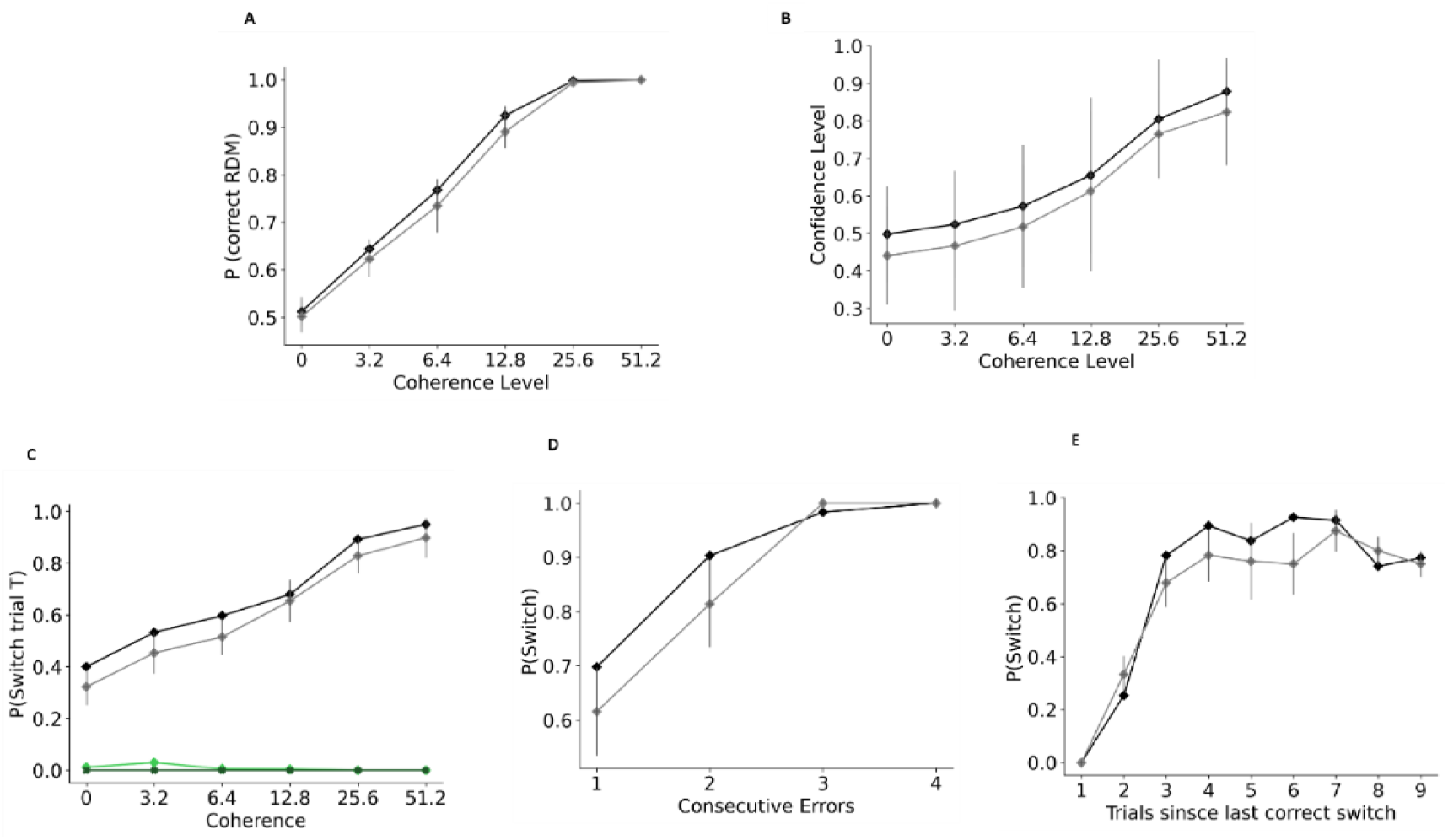
The model’s data reflect both low-level and high-level decision-making processes observed in behavioral data. **A. Probability of correct directional choice.** As coherence levels increase, so does the likelihood of correct directional choices. **B. Probability of reported confidence**. Confidence levels are strongly correlated with correct directional choices, increasing as coherence levels rise. **C. Probability of switching after receiving single negative feedback**. This probability increases with coherence levels. **D. Probability of switching after consecutive errors**. This probability rises with the number of errors. **E. Proportion of environment switches following negative feedback since the last correct switch**. This metric behaves akin to an urgency signal, initially increasing over trials before stabilizing after a few trials. In all panels, black lines represent model data, while gray lines represent behavioral data. Error bars indicate the standard error of the mean (SEM) across subjects.

Results showed that the high-level model successfully extracted three critical sources of evidence to mimic human participants’ behavior in environment switching. The model identified the importance of motion strength and extracted it from low-level model firing rates (Fig. 3C and Table S11; Eq. S22, β_1_ = 1.048 ± 0.046, *P* < 0.001), captured the impact of cumulative consecutive errors (Fig. 3D and Table S11; Eq. S22, β_1_ = 1.269 ± 0.184, *P* < 0.001) through trial-by-trial feedback, and reflected trial persistence through its prior decisions and input about the current environment (Fig. 3E and Table S11; Eq. S22, β_1_ = 0.845 ± 0.076, *P* < 0.001).

### The Model extract confidence to mimic trial by trial variability in switching decisions

As shown in Figure 3, the model effectively reproduced human switching probabilities by extracting the same information used by participants. Additionally, it accounted for trial-by-trial variability using the fine-tuned firing rates of the low-level model. A sample of trials is shown in Figure 4A, demonstrating that the model not only explains average probabilities but also captures trial-by-trial fluctuations.

**Figure 4.**
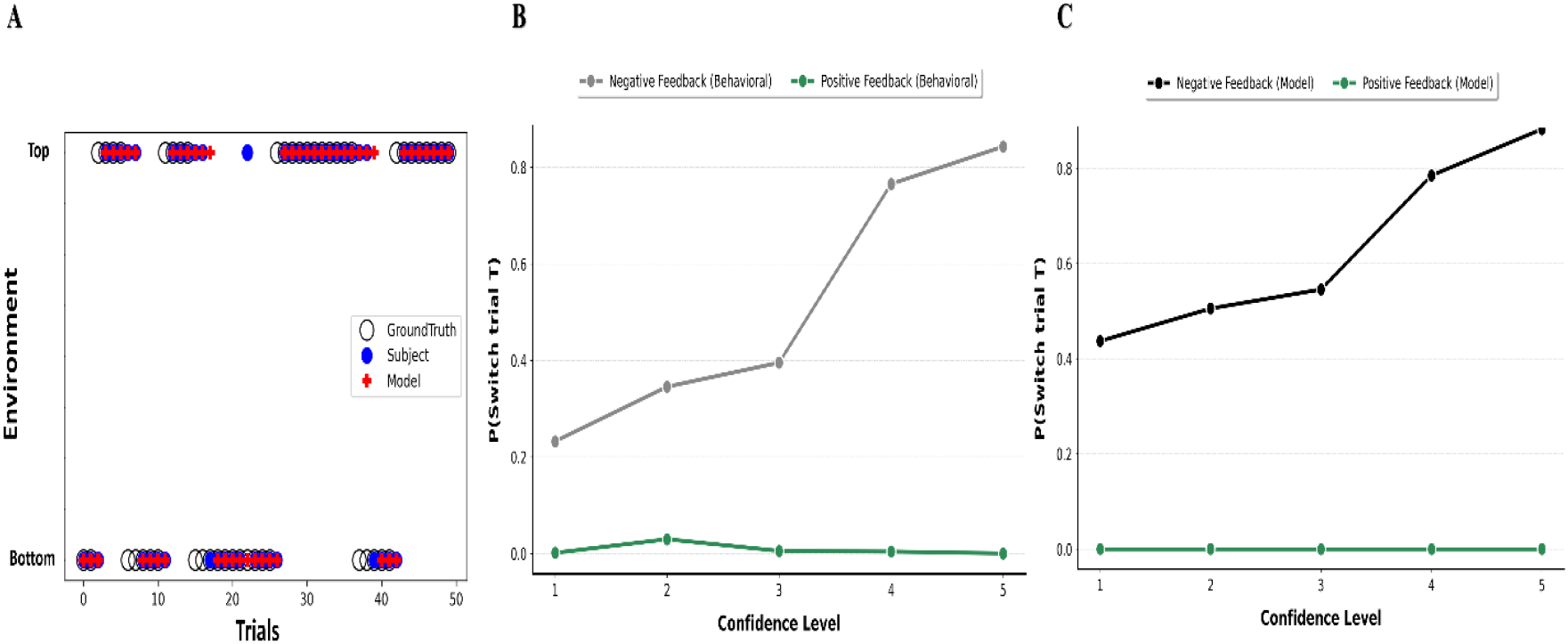
The model demonstrated the ability to infer the relationship between confidence and the probability of switching directly from signal analysis, without relying on explicit confidence ratings. **A. Trial-by-trial variability.** A schematic illustrating model choices (red crosses) and subject choices (blue points) for high-level decisions (environments) compared to ground truth (circles) over 50 test trials. **B. Subject Switching Behavior**. The probability of switching after receiving one negative feedback increased with subjects’ reported confidence levels (gray line). **C. Model Switching Behavior**. Consistent with human behavior, the model’s probability of switching after one negative feedback also increased with its internal confidence level (black line). In contrast, the probability of switching after positive feedback (green line) remained relatively stable across confidence levels.

These firing rates were fine-tuned such that the difference between the winner and loser accumulators over the 500 ms stimulus presentation conveyed participant-specific confidence on a trial-by-trial basis. To investigate whether the model captured this implicit confidence to explain trial-level variability, we examined its influence on high-level decisions within each stimulus strength level. Consistent with Purcell and Kiani (2016), switching probabilities after a single negative feedback increased with higher confidence levels (Fig. 4B). Despite lacking explicit confidence inputs, the model inferred confidence from the trial-by-trial firing rates, closely aligning with participant behavior (Fig. 4C).

### Neural Representation of Hierarchical Decision Variables (Mean Firing rate)

The LSTM model revealed evidence accumulation for switching decisions in its neural firing patterns, resembling the accumulation-to-bound mechanism proposed by Purcell and Kiani (2016) ^6^ and observed in ACC activity in monkeys ^5^. Neural responses were higher for correct trials compared to errors, and firing rates increased after two consecutive errors versus one (Fig. 5A and 5B).

**Figure 5.**
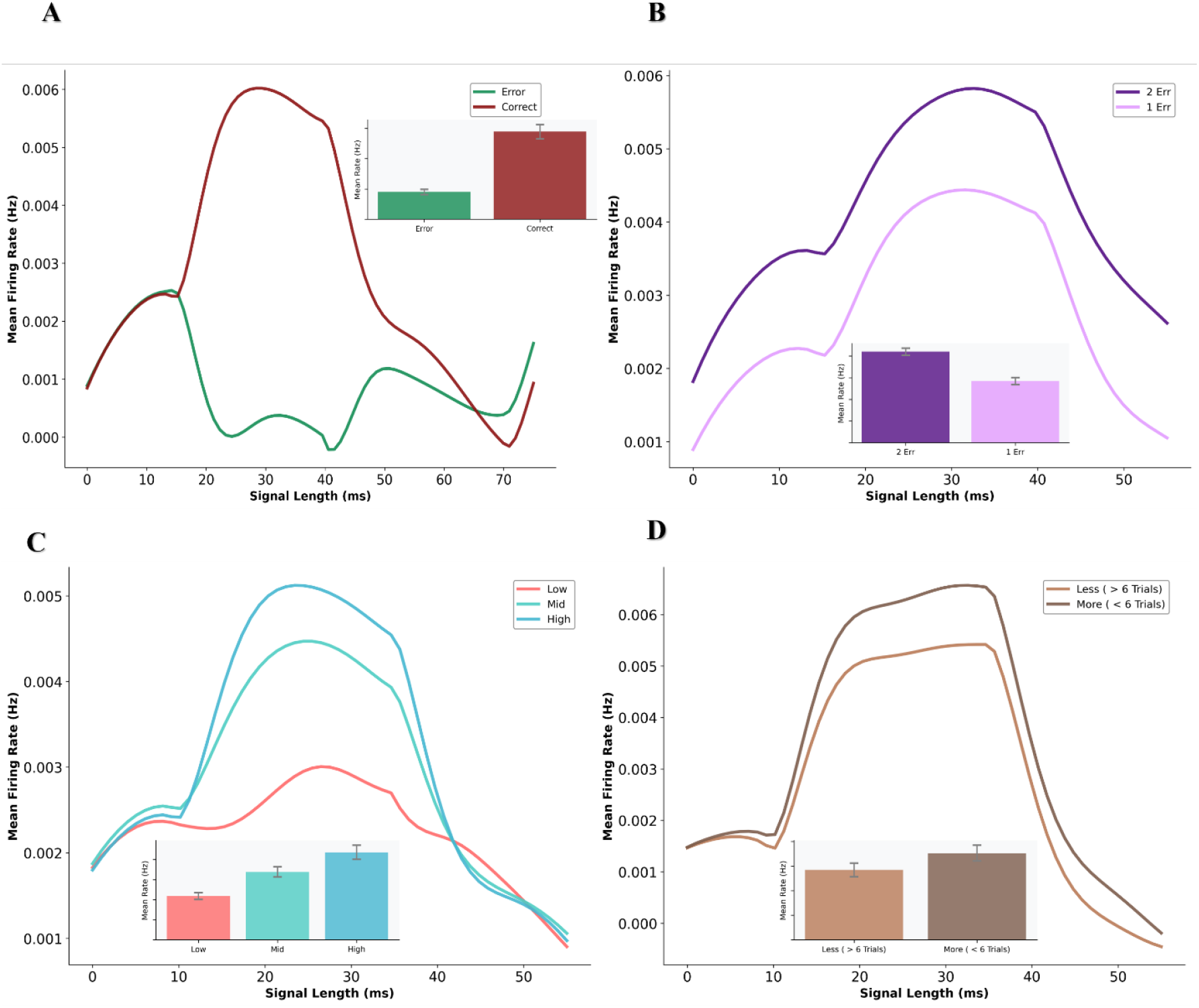
The model replicates the firing rate patterns observed in monkey ACC neurons (Refer to figure number of Saraf paper) ^5^. **A. Trial Outcome and Neural Activity.** The panel shows the average firing rates for rewarded (green) and error (red) trials. **B. Consecutive errors and Neural Activity** The panel depicts the average firing rates following one error (light purple, 1 Error) and two consecutive errors (dark purple, 2 Errors). **C. Confidence Levels and Neural Activity**. Firing rates of neurons vary across different confidence levels. Inset: Mean firing rates are modulated by both confidence level and trial outcome. (Red: Low confidence, Green: Mid confidence, Blue: High confidence). **D. Firing Rates and Urgency Signal**. Neural firing rates differ based on how long the model remained in a given environment before switching. (Light brown: Mean firing rates for switches after 1–6 trials in the same environment; Dark brown: Mean firing rates for switches after 7–12 trials). Inset: Comparison of mean firing rates in both conditions. Error bars are SEM.

We also examined the relationship between neural activity and confidence levels. Trials in which the model exhibited higher confidence in low-level decisions (Eq. S8) corresponded to elevated neural activity (Fig. 5C). Finally, neural activity reflected trial persistence, with higher average firing rates observed during longer stays in an environment (7–12 trials) compared to shorter ones (1–6 trials) (Fig. 5B). These results confirm that the model accumulates switching evidence and represents this accumulation in its neural firing rates, consistent with findings from primate studies ^5^.

### LSTM’s Internal Representation of Hierarchical Decision

As shown in Figures 3 and 4, the model not only captures the average behavior of hierarchical decision-making but also accounts for trial-by-trial variability. This demonstrates the model’s ability to effectively represent the dynamics of hierarchical decision-making. Although this outcome may not be surprising given the model’s large number of parameters, it nonetheless offers valuable insights into how such behavior is produced.

To further explore this, we examined the model’s internal representations during decision-making. Using PCA analysis, we investigated whether different hierarchical decision variables are encoded in distinct ways. Figure 6 shows that the length of the trajectory is indicative of switch probability. Specifically, in trials with a higher switch probability, neural activity traverses a larger region in latent space—taking a longer internal path when the model encounters trials closer to a switch (i.e., following more consecutive errors and with higher confidence after negative feedback). Overall, these results suggest that the model encodes the probability of a switch in the length of its internal representation trajectory.

**Figure 6.**
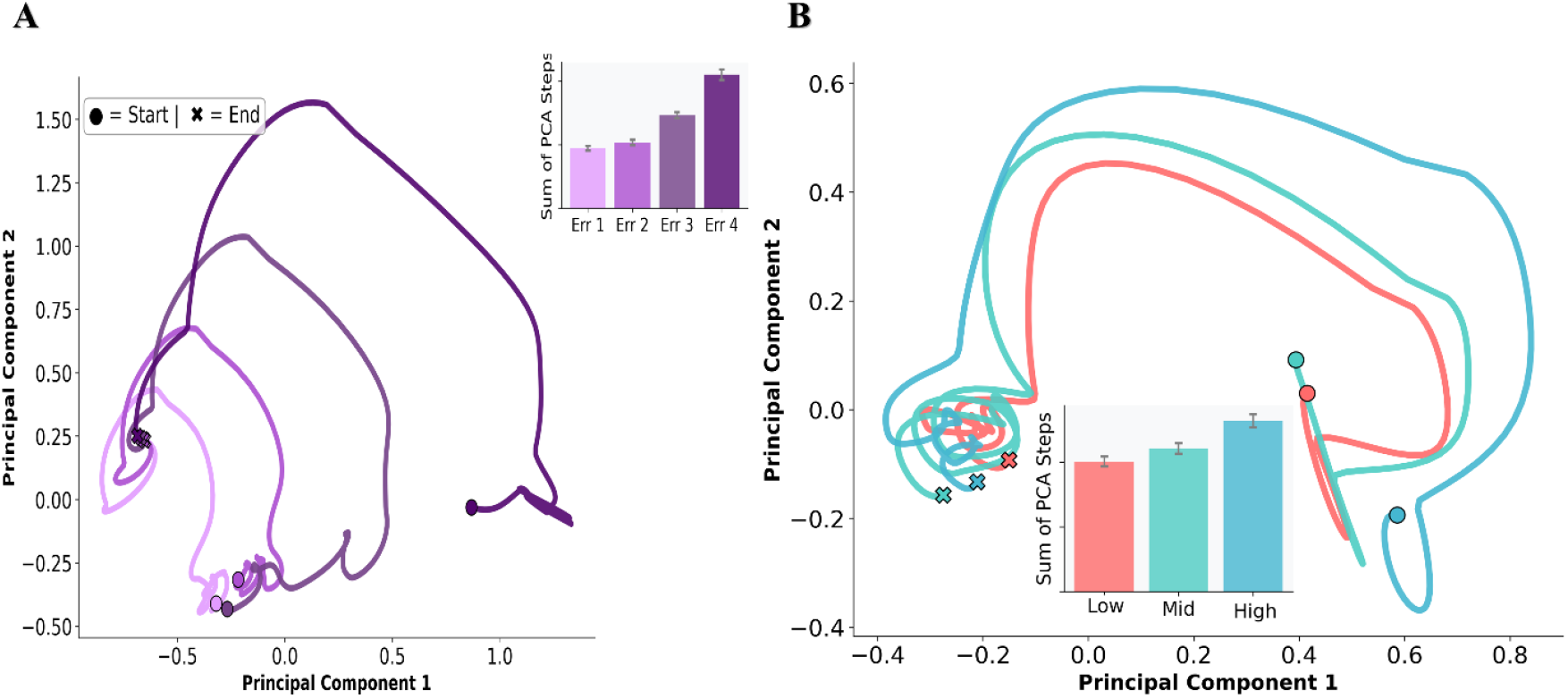
**A. Consecutive Errors**. Trajectory movement of neural activity as a function of the number of consecutive errors received. **B. Confidence Levels Following Negative Feedback**. Trajectory movement of neural activity for each confidence level when negative feedback is received. The inner bar plots show the sum of trajectory steps, indicating the total distance traversed in neural activity space. Dots indicate starting points, and crosses indicate final points.

### Feature Importance Analysis

The rich machine learning literature has provided methods to better understand black-box neural network models through a set of techniques known as explainable AI (XAI) methods. Among these, one of the most commonly used and straightforward approaches is feature sensitivity analysis, which measures how changes in input features contribute to changes in the model’s output. In our context, this allows us to quantify the importance of each input feature in determining the probability of a switch—essentially revealing how the model incorporates inputs to make its predictions. Figure 7 shows that feedback is the most critical factor influencing switch probability, followed by the winner and loser accumulators.

**Figure 7.**
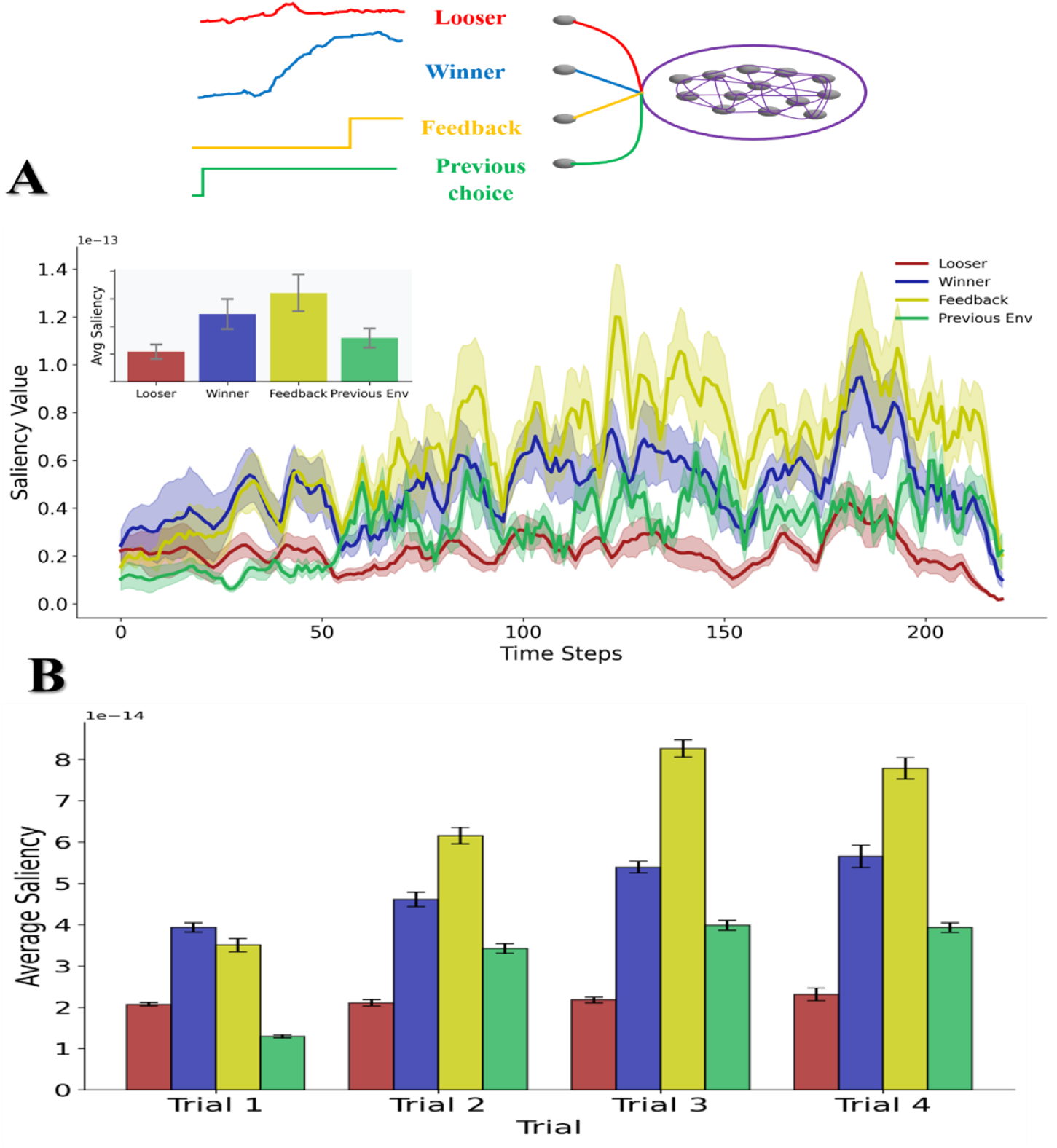
Saliency Map for Feature Importance in the Model. **A. Saliency Map Across Time Steps.** Saliency of model features at each time step, examining low-level decision signals (Loser: red, Winner: blue), feedback (yellow), and prior environment (green). **B. Recency Effect**. Averaged saliency map of model features across four consecutive trials, illustrating how feature importance evolves over time and highlights the dominance of recent feedback. Error bars are SEM.

Next, we examined the Recency Effect by analyzing the average saliency of features across four consecutive trials (corresponding to the window size). The Recency Effect refers to the phenomenon where recent trials disproportionately influence decision-making. Our results show that feedback from the most recent trial consistently exhibits the highest saliency, highlighting its dominant role in shaping the model’s predictions. Interestingly, saliency also gradually increases for feedback from earlier trials, indicating a cumulative effect over time. The winner and loser accumulators exhibit intermediate and lower saliency, respectively, further confirming that the model primarily relies on recent outcomes—particularly feedback—when predicting future switches.

Overall, these findings emphasizing the model reliance on immediate past experiences to guide decision-making.

## Discussion

Hierarchical decision-making, as the name suggests, consists of more than one stage of decision-making, where the outcome of one stage influences decisions at higher levels. In the simplest case—when the lower-level decision is a standard two-alternative forced choice (2AFC) task— there exists extensive empirical and theoretical work modeling such decisions ^22,41,48,49^. Building on this foundation, hierarchical decision-making can also be modeled. In the domain of simple perceptual tasks, the studies most closely related to ours are those by Purcell and Kiani (2016) and Sarafyazd and Jazayeri (2019) ^5,6^. They employed sequential sampling models to simulate both higher- and lower-level decisions. In our study, we propose a novel model with two key features: (1) For lower-level decision-making, we used a biologically plausible dynamic attractor network. (2) For higher-level decisions, we implemented a recurrent neural network (RNN).

Statistical models have been instrumental in shaping our understanding of decision behavior ^18,36,50–52^. However, how these models map onto neural implementation levels remains largely unclear. Biologically plausible models, such as attractor dynamic networks, may offer a framework in which decision-making can be interpreted not only computationally but also at the level of neural implementation. For this reason, we adopted this type of model for low-level decisions.

Higher-level decision-making, on the other hand, is much less explored. Both empirical and theoretical research in this domain is significantly sparser than in the case of lower-level decisions. For that reason, we opted for a computational framework that relies on as few assumptions as possible about the behavioral dynamics of the agent. Recurrent neural networks are a plausible choice for such purposes ^53–56^. We employed a powerful and overparameterized RNN model, not only to fit the data but also in the hope of reverse-engineering the model to gain insight into how it solves hierarchical decision problems. In other words, the primary value is not in the model’s ability to fit the data—which is expected—but in understanding the mechanism it develops to solve the problem.

There are now many promising methods for opening up the RNN “black box”^57^, offering pathways to gain mechanistic insight into how these models function. Several studies have analyzed RNN dynamics by constraining the rank of the connectivity matrix ^30,58,59^ or by using perturbation analysis ^60^. Many others have examined RNNs in representational or latent space ^26,61^. Therefore, although RNNs can be difficult to interpret, there are promising analytical techniques available. In our work, we employed representational analysis via PCA (Figure 6) and a relatively underutilized explainable machine learning method: sensitivity analysis (Figure 7).

Crucially, we showed that our model can replicate both behavioral and neural patterns previously reported in the literature ^1,5,6^. Behaviorally, the model exhibited non-trivial trial-by-trial variability in the probability of switching (Figure 3). Although the model was tuned to fit single-trial behavior (e.g., accuracy and confidence), computing the trial-by-trial probability of switching is non-trivial—suggesting that the model captures more than just average behavior and reflects internal variability as well.

Furthermore, our model offers a flexible framework for investigating the mechanisms underlying confidence. Although the present study did not explicitly focus on this aspect, the implicit emergence of confidence from the network’s dynamics presents a unique opportunity to explore its computational and functional contributions to hierarchical decision-making. This is particularly important given that confidence has been extensively studied ^41,62–64^ and is known to serve as a bridge for credit assignment across different levels of a decision hierarchy ^1,6^.

Neurally, the average activity of our model exhibited qualitative similarities to previously reported data from the anterior cingulate cortex (ACC) showing ramping activity in trials with a higher probability of switching. In the case of low-level decision models, prior studies have demonstrated how the attractor model corresponds to specific brain areas (e.g., ^37^). However, this is not yet the case for our high-level RNN model. While the RNN may loosely correspond to activity in the ACC, further empirical and theoretical work is needed to clarify this relationship.

To open up the RNN black box, we first used representational analysis via PCA. This revealed that trials with a higher likelihood of switching spanned a broader latent space than other trials. This effect may reflect a latent-space signature of evidence accumulation, similar to population-level neural dynamics (Figure 5). The result aligns with recent findings showing evidence accumulation in contexts beyond the random dot motion task ^65^.

Additionally, we applied a set of explainable AI methods to analyze the model’s sensitivity to its inputs. These techniques aim to identify which input features the neural network relies on most heavily when making predictions. We quantified the sensitivity of the RNN inputs and found that feedback was the most influential factor in the model’s decision to switch environments. Further techniques—such as LIME ^66^ and SHAP ^67^—could be employed to better understand feature interactions.

Overall, this approach enables the use of powerful neural network models while still striving for interpretability and insight into the mechanisms underlying complex decision-making processes.

## Acknowledgments

JE Was supported by the European Research Council (ERC) under the European Union’s Horizon 2020 research and innovation programme (819040 - acronym: rid-O).

## Notes

### Competing Interest Statement

The authors have declared no competing interest.

